# Exercise training adaptations in liver glycogen and glycerolipids require hepatic AMP-activated protein kinase in mice

**DOI:** 10.1101/2023.09.01.555935

**Authors:** Curtis C. Hughey, Deanna P. Bracy, Ferrol I. Rome, Mickael Goelzer, E. Patrick Donahue, Benoit Viollet, Marc Foretz, David H. Wasserman

## Abstract

Regular exercise elicits adaptations in glucose and lipid metabolism that allow the body to meet energy demands of subsequent exercise bouts more effectively and mitigate metabolic diseases including fatty liver. Energy discharged during the acute exercise bouts that comprise exercise training may be a catalyst for liver adaptations. During acute exercise, liver glycogenolysis and gluconeogenesis are accelerated to supply glucose to working muscle. Lower liver energy state imposed by gluconeogenesis and related pathways activates AMP-activated protein kinase (AMPK), which conserves ATP partly by promoting lipid oxidation. This study tested the hypothesis that AMPK is necessary for liver glucose and lipid adaptations to training. Liver-specific AMPKα1α2 knockout (LAKO) and wild type (WT) mice completed sedentary and exercise training protocols. Liver nutrient fluxes were quantified at rest or during acute exercise following training. Liver metabolites and molecular regulators of metabolism were assessed. Training increased liver glycogen in WT mice, but not in LAKO mice. The inability to increase glycogen led to lower glycogenolysis, glucose production, and circulating glucose during acute exercise in trained LAKO mice. Deletion of AMPKα1α2 attenuated training-induced declines in liver diacylglycerides. In particular, training lowered the concentration of unsaturated and elongated fatty acids comprising diacylglycerides in WT mice, but not in LAKO mice. Training increased liver triacylglycerides and the desaturation and elongation of fatty acids in triacylglycerides of LAKO mice. These lipid responses were independent of differences in tricarboxylic acid cycle fluxes. In conclusion, AMPK is required for liver training adaptations that are critical to glucose and lipid metabolism.

**NEW & NOTEWORTHY:** This study shows that the energy sensor and transducer, AMP-activated protein kinase, is necessary for an exercise training-induced: i) increase in liver glycogen that is necessary for accelerated glycogenolysis during exercise, ii) decrease in liver glycerolipids independent of TCA cycle flux, and iii) decline in the desaturation and elongation of fatty acids comprising liver diacylglycerides. The mechanisms defined in these studies have implications for use of regular exercise or AMPK-activators in patients with fatty liver.

## INTRODUCTION

Exercise accelerates liver glucose production to match the higher rate of muscle glucose uptake (1, 2). The increased liver glucose production is accomplished by a rise in both liver glycogenolysis and gluconeogenesis (1–6). *De novo* synthesis of glucose via gluconeogenesis is energetically costly and contributes to the elevated hydrolysis of ATP to ADP and AMP during exercise (7). To mitigate this energy discharge and support the increased gluconeogenic flux, fatty acid oxidation within the liver is stimulated (5, 6, 8). Thus, acute exercise requires that the liver transform the potential energy in fatty acids to the common energy currency of ATP (9). Importantly, the repeated provocation of energy discharge by the individual exercise bouts that comprise training is associated with liver glucose and lipid metabolic adaptations that allow the body to better meet the energy demands of subsequent acute exercise. Training increases liver glycogen, elevates capacity and flexibility to dispose of lipids in mitochondrial oxidative metabolism pathways, and decreases lipogenesis (8–16). These adaptations in glucose and lipid metabolism may also support the efficacy of training to mitigate liver steatosis and maintain glycemic control in obesity and diabetes (7–11). While the metabolic adaptations of the liver to training and their public health implications have been investigated, the extent to which the repeated energy discharge of exercise mediates liver adaptations to training and the key underlying molecular mechanisms of action remain to be clearly defined.

Prior work has indicated that intracellular sensing and responses to protect against a more severe fall in energy state are critically reliant on AMP-activated protein kinase (AMPK) (17). AMPK is a heterotrimeric serine/threonine kinase consisting of a catalytic α subunit and regulatory β and γ subunits (18). The α subunit exists as one of two isoforms (α1 and α2) in hepatocytes and is the site of kinase activity that is stimulated by a lowered energetic state (19, 20). More specifically, ATP and ADP hydrolysis results in AMP accumulation which binds to the γ subunit resulting in a conformational change that promotes activation by the phosphorylation of threonine residue 172 of the α subunits (18). Once activated, AMPK-mediated phosphorylation events control cellular metabolism in a manner that favors restoration of ATP (21). These include promotion of carbohydrate and lipid catabolic pathways (21). In addition to regulation by energy state, the β subunit of AMPK contains a carbohydrate-binding module that binds to glycogen, which is suggested to inhibit AMPK (22). Importantly, the activation of AMPK during acute exercise is closely linked to a decline in ATP and glycogen (5, 23).

Given that AMPK is a sensor and transducer of energy discharge elicited during acute exercise, this study tested the hypothesis that hepatic AMPK is necessary for key liver glucose and lipid adaptations to repeated exercise bouts (i.e., exercise training). This was accomplished by studying mice with a liver-specific knockout of AMPK α1 and α2 subunits (LAKO) and wild type (WT) littermates that were untrained or underwent exercise training. *In vivo* isotope infusions combined with ^2^H/^13^C metabolic flux analysis quantified liver glucose and mitochondrial oxidative fluxes at rest and during exercise. In addition, the importance of AMPK on key metabolite concentrations and the expression of molecular regulators of liver nutrient metabolism were examined. The results of this study show that liver glycogen and glycerolipid adaptations to regular exercise are dependent on hepatic AMPK.

## MATERIALS AND METHODS

### Mouse models and husbandry

The Vanderbilt University Animal Care and Use Committee approved all mouse procedures. Mice with a double hepatic knockout of AMPK α1 and α2 subunits were created by breeding mice expressing Cre recombinase under control of the albumin promoter (B6.Cg-Tg(Alb-cre)21Mgn/J) with C57BL6/J mice harboring AMPK α1 and α2 alleles flanked by loxP sites (5). Floxed AMPKα1α2 littermates of LAKO mice that did not have transgenic Cre recombinase were used as WT control mice. Deletion of AMPK α1 and α2 subunits in the liver of mice was confirmed by both PCR genotyping and immunoblotting. Male mice receiving Laboratory Rodent Diet (LabDiet^®^ 5001, St. Louis, MO, USA) and water ad libitum were studied. Mice were housed on natural soft cellulose bedding (BioFresh™ Comfort Bedding, Ferndale, WA, USA) under temperature- and humidity-controlled conditions and a 12-hour light/dark cycle.

### Exercise stress test

Mice completed two exercise stress tests. The first was performed at 29 weeks of age and completed 24 hours prior to the start of a 6-week exercise training protocol. The second was performed 24 hours following the completion of the 6-week training protocol. The stress test protocols were completed to define the maximal running speed of mice as previously described (5, 24). Briefly, mice were acclimated to running on an enclosed single lane treadmill (Columbus Instruments, Columbus, OH, US) by performing two 10-min exercise bouts at 10 m/min (0% incline) 24 and 48 hours prior to the first exercise stress test and the second stress test in untrained mice. For the exercise stress test, mice were placed in the same enclosed single lane treadmill. Following a 10-min sedentary period, mice initiated running at 10 m·min^-1^ (0% incline). The treadmill belt speed was increased by 4 m·min^-1^ every 3 min until exhaustion. Exhaustion was defined as the point in which the mouse remained on the shock grid at the posterior of the treadmill for more than five continuous seconds.

### Exercise training protocol

Fig. 1A provides a schematic representation of the training protocol and timeline. Mice completed a 60 min treadmill running bout, five days a week for six weeks. The treadmill running speed was 45% of the mouse’s initial maximal running speed during the first week of training and increased progressively by 5% of the maximal running speed each week such that the mice were running at 70% of their initial maximal running speed during the sixth week of training. Untrained mice were placed into an enclosed treadmill in which the treadmill belt did not move. Like the trained mice, the intervention in untrained mice was five days per week, 60 min per day for six weeks.

**FIGURE 1.**
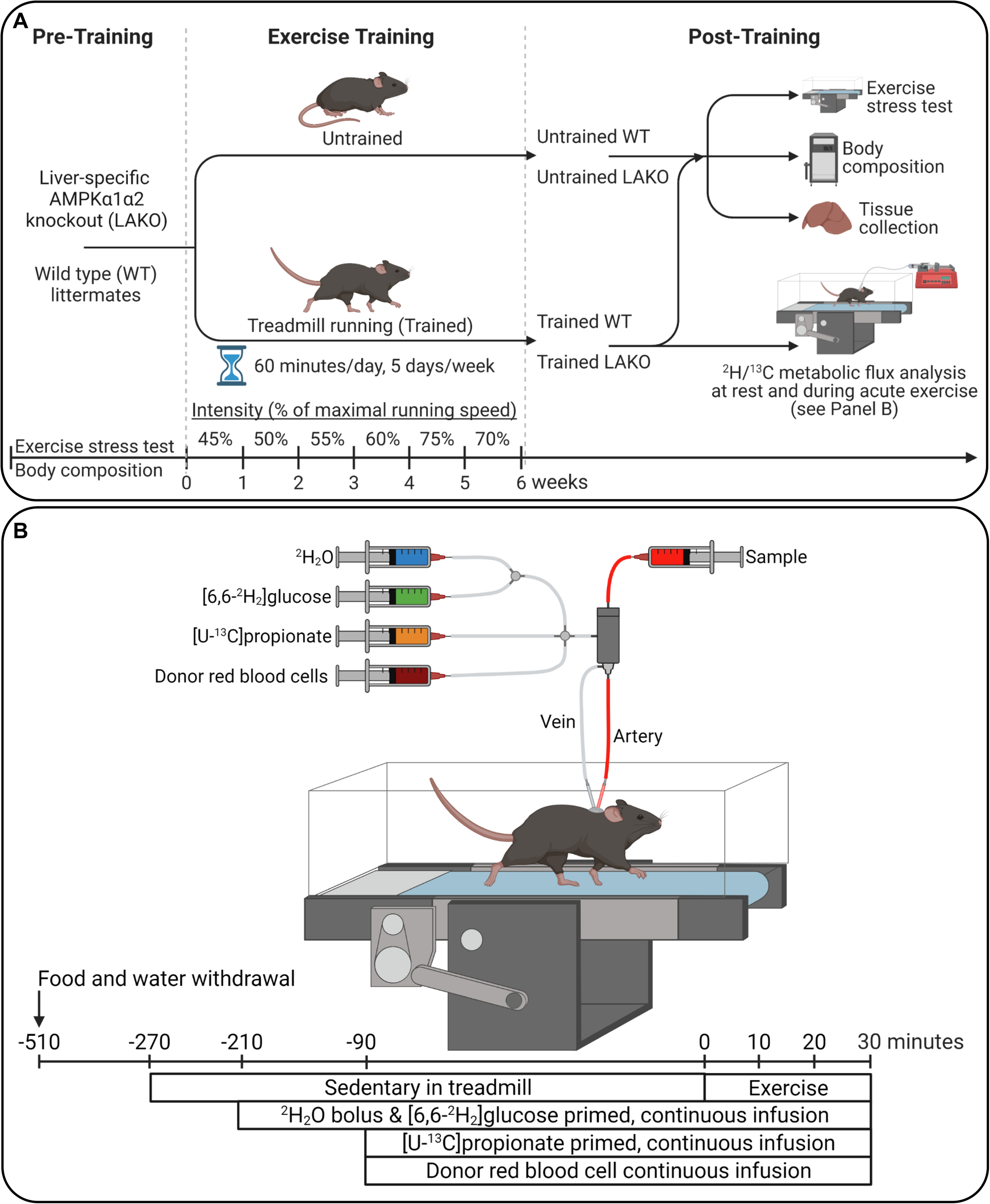
Schematic representation of experimental timeline and procedures. **(A)** Mice with a liver-specific deletion of AMPK α1 and α2 subunits (Liver AMPK KO) and wild type (WT) littermates were randomly assigned to untrained and exercise training protocols. The training protocol consisted of treadmill running 60 minutes a day, five days per week for 6 weeks. Pre- and post- training analyses included exercise stress tests to evaluate maximal running speed and body composition. A cohort of trained mice underwent vascular surgeries for isotope infusion experiments and liver nutrient flux quantification. Tissues were collected from mice upon completion of *in vivo* experiments for molecular analysis. **(B)** Stable isotope infusions during acute treadmill running bout were performed in trained mice 10 days following carotid arterial and jugular catheter implantation surgeries. At 210 minutes prior to the treadmill running bout (5 hours of fasting), a ^2^H_2_O bolus was administered into the venous circulation to enrich total body water at 4.5%. A [6,6-^2^H_2_]glucose prime was infused followed by a continuous infusion was initiated with the ^2^H_2_O bolus. Ninety minutes before the onset of exercise (7 hours of fasting), a primed, continuous infusion of [U-^13^C]propionate was started. Donor red blood cells were administered at a constant rate to prevent a decline in hematocrit due to arterial sampling. Arterial samples were obtained prior to stable isotope infusion as well as during 30-minute exercise bout for ^2^H/^13^C metabolic flux analysis.

### Body composition

The first body composition measurements were obtained via an mq10 nuclear magnetic resonance analyzer (Bruker Corporation, Billerica, MA, US) in 29-week-old mice prior to the first acclimation exercise bout of the stress test procedures. The second measurement was determined in 35-week-old mice prior to the final 60-min treadmill running bout.

### Surgical procedures

Twenty-four hours following the final exercise stress test, a cohort of trained mice underwent vascular catheter implantation procedures (25). Briefly, catheters were implanted in the jugular vein and carotid artery for infusions and sampling, respectively. The exteriorized ends of the implanted catheters were flushed with 200 U·ml^-1^ heparinized saline and 5 mg·ml^-1^ ampicillin and sealed with stainless-steel plugs. Following surgery, mice were housed individually and provided 10 days of post-operative recovery prior to stable isotope infusions studies being performed. To prevent detraining during the 10-day surgical recovery period, mice completed five, 60-minute treadmill running bouts at 45% of their maximal running speed determined from the second exercise stress test on days 3 to 8 following the surgical procedures.

### Stable isotope infusions

Stable isotope infusions were performed at rest and during acute exercise. The purpose of acute exercise was to study nutrient fluxes during a stimulus that is likely to expose adaptations to training should they exist. All mice were within 10% of pre-surgical weight prior to stable isotope infusions. Beginning in the first hour of the light cycle both food and water were withdrawn for the entirety of the experiment (9 hours). Four hours into the fast, mice were placed in an enclosed single lane treadmill and the exteriorized catheters were connected to infusion syringes. Following a one-hour acclimation period, an 80 μl arterial blood sample was obtained to determine natural isotopic enrichment of plasma glucose. Immediately after this sample acquisition, a stable isotope infusion protocol was initiated to allow for the quantification of endogenous glucose production and associated oxidative fluxes as previously performed (5, 6, 26–30). Briefly, a ^2^H_2_O (99.9%)-saline bolus containing [6,6-^2^H_2_]glucose (99%) was infused over 25 minutes to concurrently label the H_2_O pool and provide a [6,6-^2^H_2_]glucose prime (440 μmol·kg^-1^). An independent, continuous infusion of [6,6-^2^H_2_]glucose (4.4 μmol·kg^-1^·min^-1^) was started following the ^2^H_2_O-saline bolus and [6,6-^2^H_2_]glucose prime. A primed (1.1 mmol·kg^-1^), continuous (0.055 mmol·kg^-1^·min^-1^) intravenous infusion of [U-^13^C]propionate (99%, sodium salt) was started two hours after the ^2^H_2_O bolus and [6,6-^2^H_2_]glucose prime. Four arterial blood samples (100 μl) were obtained 90-120 minutes following the [U-^13^C]propionate bolus (time = 0-30 minutes of treadmill running) to determine arterial blood glucose using an Accu-Chek® glucometer (Roche Diagnostics, Indianapolis, IN, USA) and to perform ^2^H/^13^C metabolic flux analysis. The sample taken at 90 minutes following the [U-^13^C]propionate bolus (time = 0 minutes) was obtained while mice were in a sedentary state on a stationary treadmill. Samples taken 100-120 minutes following the [U-^13^C]propionate bolus (time = 10-30 minutes of exercise) were obtained while mice were completing an acute treadmill running bout at 35% of their post-training maximal running speed. Plasma samples were stored at −20°C for subsequent isotopic analysis. Donor erythrocytes were infused throughout the experiment to maintain hematocrit constant. Mice were removed from the treadmill and sacrificed by cervical dislocation immediately after the final sample was taken. Liver tissues were rapidly excised, freeze-clamped in liquid nitrogen, and stored at −80°C until they were used for metabolite analyses and immunoblotting. A schematic of the isotope infusion protocol is provided in Fig. 1B.

### Glucose derivatization and gas-chromatography-mass spectrometry (GC-MS) analysis

Approximately 40 μl of plasma obtained prior to the stable isotope infusion and at the 0-, 10-, 20-, and 30-minute time points of the treadmill running bout were used for di-*O*-isopropylidene propionate, aldonitrile pentapropionate, and methyloxime pentapropionate derivatives of glucose (5). GC-MS analysis was performed and uncorrected mass isotopomer distributions (MIDs) for six fragment ions were determined as previously described (5, 28, 29).

### 2H/13C Metabolic flux analysis

The details of the *in vivo* metabolic flux analysis methodology employed in these studies have been described previously (27). In summary, a reaction network was constructed using Isotopomer Network Compartmental Analysis (INCA) software (31). The reaction network defined the carbon and hydrogen transitions for hepatic glucose production and associated oxidative metabolism reactions. The flux through each network reaction was determined relative to citrate synthase flux (V_CS_) by minimizing the sum of squared residuals between simulated and experimentally determined MIDs of the six fragment ions previously described (27, 32). Flux estimates were repeated 50 times from random initial values. Goodness of fit was assessed by a chi-square test (*p* = 0.0*5*) with 34 degrees of freedom. Confidence intervals of 95% were determined as previously described (27, 32). Mouse body weights and the [6,6-^2^H_2_]glucose infusion rate were used to determine absolute values.

### Tissue metabolite analyses

Glycogen (33) adenine nucleotides (34), and lipids (35) were measured in livers as described previously. Energy charge was calculated as ([ATP]+0.5[ADP])/([ATP]+[ADP]+[AMP]) (36). Fatty acyl chain desaturation indexes were determined from the palmitoleate (C16:1)-to-palmitate (C16:0) ratio and the oleate (C18:1)-to-stearate (C18:0) ratio. The fatty acyl chain elongation index was calculated as [(C18:0+C18:1)/C16:0] (37). These assays were performed in two cohorts of mice. The first being livers from trained mice following the completion of 30 minutes of acute treadmill running and the stable isotope infusion protocol. The second cohort consisted of untrained or trained, non-catheterized mice. On the day of the study the second cohort underwent a protocol designed to match the protocol for cohort one. The difference was that mice in cohort one were exercised while mice in cohort 2 were placed in an enclosed, stationary single lane treadmill (Columbus Instruments, Columbus, OH) four hours into a fast and sacrificed at 8.5 hours of fasting (equivalent to the 0 min timepoint of the stable isotope infusion studies). This protocol was designed to match the timing of cohort one so that liver metabolites determined both prior to and following acute exercise could be compared.

### Immunoblotting

Liver homogenates were prepared as previously outlined (5, 28, 29). Liver proteins (15 μg) were separated via electrophoresis on a NuPAGE 4-12% Bis-Tris gel (Invitrogen, Carlsbad, CA, USA) and subsequently transferred to a PVDF membrane. The primary and secondary antibodies used for immunoblotting are provided in Supplemental Table S1. The antibody-probed PVDF membranes were treated with chemiluminescent substrate (ThermoFisherScientific, Waltham, MA, USA) and images acquired with a ChemiDoc™ Imaging system and Image Lab™ software (Bio-Rad, Hercules, CA, USA). Total protein was measured via BLOT-FastStain (G-Bioscience, St. Louis, MO, USA) and used as a loading control. Densitometry was completed using ImageJ software.

### Statistical analyses

GraphPad Prism software (GraphPad Software LLC., San Diego, CA) was used to perform one sample t test, Student’s t test, two-way ANOVAs followed by Tukey’s post hoc tests, and repeated-measures ANOVAs followed by Sidak’s post hoc tests as appropriate to detect statistical differences (p<0.05). All data are reported as mean ± SEM.

## RESULTS

### Exercise training increases maximal running speed in WT and liver-specific AMPK KO mice

Initial experiments assessed the maximal running speed of LAKO mice and WT littermates using an exercise stress test. The results of this test were used to determine the running speed protocols during training. Prior to the onset of the 6-week training protocol, WT and LAKO mice had comparable maximal running speeds (Table 1). Training increased the maximal running speed in both WT and LAKO mice (Table 1). Body weight and composition were similar between WT and LAKO mice prior to and following training protocols (Table 1). Thus, loss of liver AMPK did not negatively impact exercise capacity or body composition. Moreover, the training-induced increase in maximal exercise capacity is independent of liver AMPK.

**Table 1.** Biometric characteristics of untrained and exercise-trained mice lacking liver AMPKα1α2. Maximal running speed (m·min^-1^) in wild type (WT) and liver-specific AMPKα1α2 knockout (LAKO) mice prior to and following 6 weeks of training (n = 9-17 mice per group). Body weight (g), Fat Mass (%), and Lean Mass (%) in WT and LAKO mice prior to and following a 6-week training protocol (n = 9-16 mice per group). Data are mean ± SEM. **p<0.01 vs. untrained WT mice following 6-weeks of training. *p<0.05 vs. untrained LAKO mice following 6-weeks of training.

|  | Untrained groups before 6-week protocol |  | Trained groups before 6-week protocol |  | Untrained groups after 6-week protocol |  | Trained groups after 6-week protocol |  |
| --- | --- | --- | --- | --- | --- | --- | --- | --- |
| Running Characteristics | WT | LAKO | WT | LAKO | WT | LAKO | WT | LAKO |
| Maximal Running Speed (m·min<sup>-1</sup>) | 34.0 ± 3.0 | 29.6 ± 1.8 | 34.0 ± 1.1 | 32.8 ± 1.7 | 30.4 ± 2.9 | 31.8 ± 2.3 | 41.8 ± 1.7** | 42.3 ± 2.3* |
| Body Composition | WT | LAKO | WT | LAKO | WT | LAKO | WT | LAKO |
| Body Weight (g) | 29.6 ± 0.4 | 29.7 ± 0.9 | 30.7 ± 0.4 | 31.0 ± 0.7 | 29.8 ± 0.6 | 29.8 ± 1.0 | 29.9 ± 0.4 | 30.6 ± 0.4 |
| Fat Mass (%) | 8.0 ± 0.8 | 7.2 ± 0.5 | 9.9 ± 0.6 | 9.9 ± 1.0 | 8.6 ± 0.6 | 8.9 ± 0.7 | 10.7 ± 0.4 | 10.5 ± 0.8 |
| Lean Mass (%) | 71.3 ± 0.6 | 71.7 ± 0.7 | 69.7 ± 0.5 | 69.7 ± 1.0 | 70.6 ± 0.6 | 70.0 ± 0.5 | 68.9 ± 0.5 | 68.8 ± 0.9 |

### Hepatic AMPK protects against exercise training-induced energetic stress

While training increased muscular performance to a similar extent in both genotypes, a lack of liver AMPKα1α2 led to differential responses in liver energy state (Fig. 2A-G). The six-week training protocol did not influence the phospho-AMPKα^Thr172^ (pAMPK)-to-AMPK ratio (Fig. 2A). Liver ATP was comparable between untrained WT and LAKO mice (Fig. 2A). However, an interaction effect was observed that led to higher liver ATP in trained WT compared to trained LAKO mice (Fig. 2B). Liver AMP (Fig. 2D), the AMP-to-ATP ratio (Fig. 2F), and energy charge (Fig. 2G) also showed interaction effects. Thus, hepatic AMPK is necessary for training to better maintain liver energy state under fasting, sedentary conditions.

**FIGURE 2.**
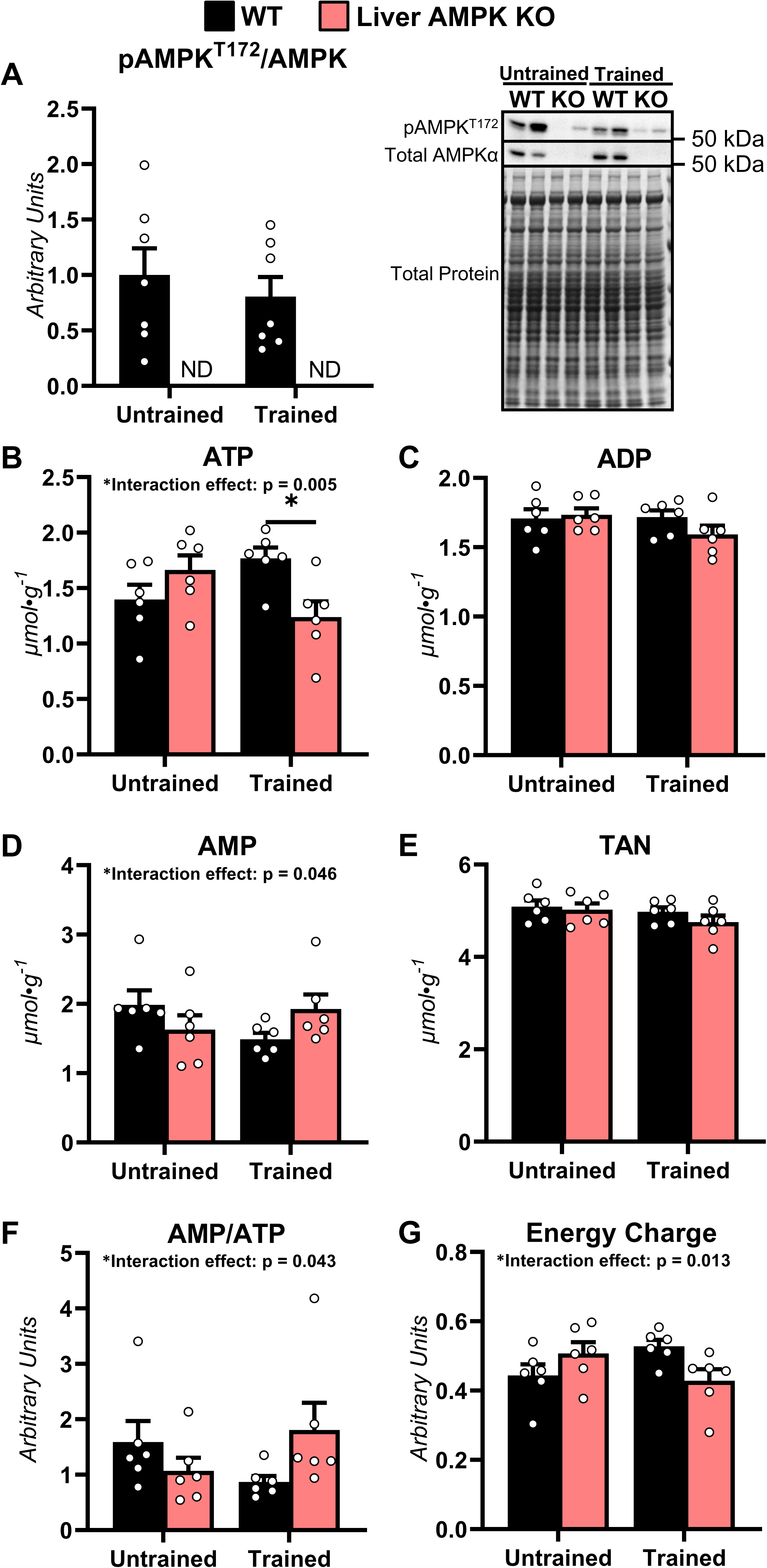
Liver energy homeostasis in untrained and exercise-trained mice lacking hepatic AMPK. Livers from untrained and exercise-trained mice with liver-specific deletions of AMPK α1 and α2 subunits (Liver AMPK KO) and wild type (WT) littermates were obtained from non-catheterized mice on a stationary treadmill at the 0-minute time point (8.5-hour fast). **(A)** Liver phosphorylated AMPKα-to-total AMPK (pAMPK^T172^/AMPK) as determined by immunoblotting and representative immunoblots. Liver **(B)** ATP (μmol·g^-1^), **(C)** ADP (μmol·g^-1^), **(D)** AMP (μmol·g^-1^), and **(E)** total adenine nucleotide pool (μmol·g^-1^; TAN = ATP + ADP + AMP). Liver **(F)** AMP/ATP and **(G)** energy charge (arbitrary units; [ATP+0.5ADP]/[TAN]). n = 6-7 per group. Data are mean ± SEM. *p<0.05 by two-way ANOVA followed by Tukey’s post hoc tests.

### Molecular regulators of mitochondrial metabolism are decreased in livers of LAKO mice

Mitochondrial oxidative metabolism is a primary controller of energy homeostasis. The expression of mitochondrial proteins was evaluated (Fig. 3A-F). Liver respiratory chain complexes III and IV were lower in untrained LAKO mice compared to untrained WT mice (Fig. 3A and D). Training increased liver respiratory chain complex III in WT mice, however, the training response was absent in LAKO mice (Fig. 3A and D). Respiratory chain complexes I and IV were also higher in livers of trained WT mice compared to trained LAKO mice (Fig. 3A and D). Liver mitochondrial pyruvate carrier 1 (MPC1), but not mitochondrial pyruvate carrier 2 (MPC2), was reduced by loss of AMPKα1α2 (Fig. 3B and D). Training modestly increased ketogenic enzyme, 3-hydroxymethylglutaryl-CoA synthase 2 (HMGCS2) (Fig. 3C and D). These results show that liver mitochondrial proteins, particularly those involved in electron transport, are decreased in LAKO mice compared to WT mice. Moreover, the differential expression of liver respiratory chain complexes between WT and LAKO mice was amplified by training.

**FIGURE 3.**
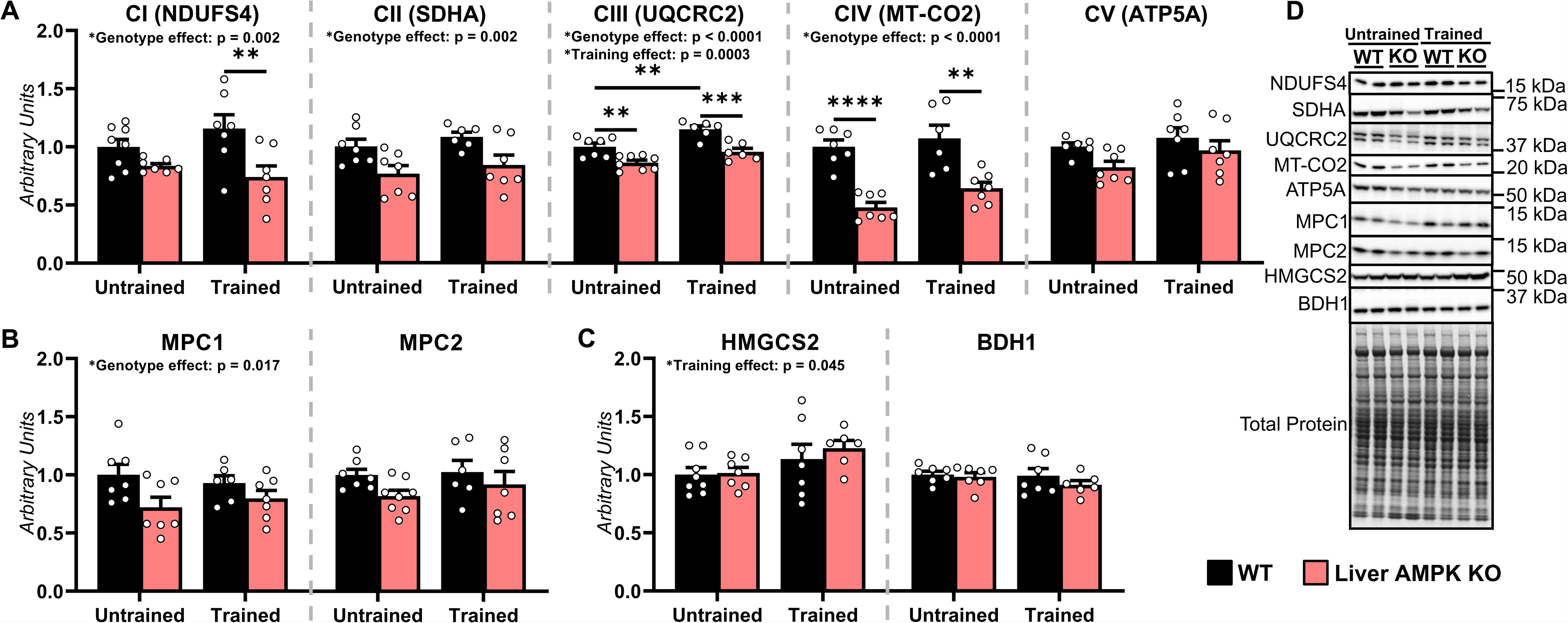
Regulators of mitochondrial oxidative metabolism in untrained and exercise-trained mice lacking hepatic AMPK. Livers from untrained and trained mice with liver-specific deletions of AMPK α1 and α2 subunits (Liver AMPK KO) and wild type (WT) littermates were obtained from non-catheterized mice on a stationary treadmill at the 0-minute time point (8.5-hour fast). Liver **(A)** respiratory chain complexes 1-5 (CI-CIV), **(B)** mitochondrial pyruvate carrier 1 and 2 (MPC1 and 2) and **(C)** 3-hydroxymethylglutaryl-CoA synthase 2 (HMGCS2) and β-hydroxybutyrate dehydrogenase (BDH1) as determined by immunoblotting with **(D)** representative immunoblots. n = 6-8 per group. Data are mean ± SEM. **p<0.01, ***p<0.001, and ****p<0.0001 by two-way ANOVA followed by Tukey’s post hoc tests.

### Changes in liver glycerolipid pool size and fatty acyl chain composition following exercise training are AMPK-dependent

Next, we assessed molecular regulators of lipid synthesis (Fig 4A). The phospho-acetyl-CoA carboxylase (pACC)-to-ACC ratio was lower in livers of LAKO mice compared to their WT littermates under both untrained and trained conditions (Fig. 4A). Liver fatty acid synthase (FASN) was comparable under all conditions, while a genotype effect showed stearoyl-CoA desaturase 1 (SCD1) was higher in LAKO mice (Fig. 4A). Liver diacylglyceride (DAG), triacylglyceride (TAG), and phospholipid concentrations in untrained WT mice were similar to that in untrained LAKO mice (Fig. 4B-D). Training reduced DAGs by ∼45% in WT mice. This decrease in DAGs was blunted in LAKO mice (Fig. 4B). An interaction effect of AMPK deletion and training exposed higher liver TAGs in trained LAKO mice compared to trained WT mice (Fig. 4C). Training reduced liver phospholipids in WT mice, but not in LAKO mice (Fig. 4D).

**FIGURE 4.**
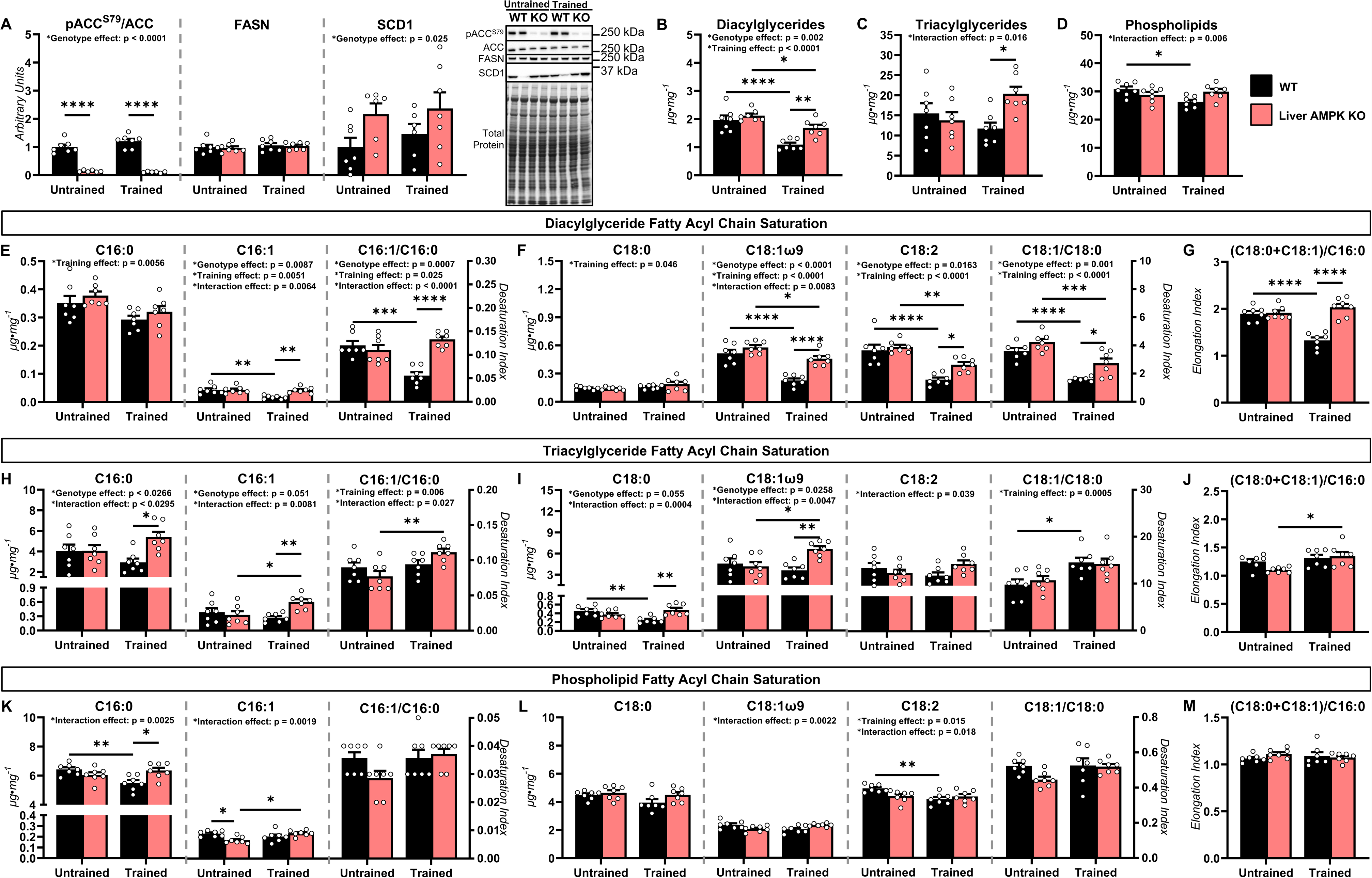
Lipid metabolites and molecular regulators in untrained and exercise-trained mice lacking hepatic AMPK. Livers from untrained and trained mice exhibiting liver-specific deletions of AMPK α1 and α2 subunits (Liver AMPK KO) and wild type (WT) littermates were obtained from non-catheterized mice on a stationary treadmill at the 0-minute time point (8.5-hour fast). **(A)** Liver phosphorylated acetyl-CoA carboxylase-to-total acetyl-CoA carboxylase ratio (pACC^S79^/ACC), fatty acid synthase (FASN), and stearoyl-CoA desaturase 1 (SCD1) as determined by immunoblotting and representative immunoblots. **(B)** Liver diacylglyceride concentration (µg·mg^-1^; DAG). **(C)** Liver triacylglyceride concentration (µg·mg^-1^; TAG). **(D)** Liver phospholipid concentration (µg·mg^-1^; PL). **(E)** C16:0 and C16:1 fatty acids (µg·mg^-1^) in liver DAGs and the C16:1-to-C16:0 ratio. **(F)** C18:0, C18:1, and C18:2 fatty acids in liver DAGs (µg·mg^-1^) and the C18:1-to-C18:0 ratio. **(G)** Elongation index [(C18:0+C18:1)/C16:0] of fatty acyl chains in liver DAGs. **(H)** C16:0 and C16:1 fatty acids in liver TAGs (µg·mg^-1^) and the C16:1-to-C16:0 ratio. **(I)** C18:0, C18:1, and 18:2 fatty acids in liver TAGs and the C18:1-to-C18:0 ratio. **(J)** Elongation index [(C18:0+C18:1)/C16:0] of fatty acyl chains in liver TAGs. **(K)** C16:0 and C16:1 fatty acids in liver PLs (µg·mg^-1^) and the C16:1-to-C16:0 ratio. **(L)** C18:0, C18:1, and C18:2 fatty acids in liver PLs (µg·mg^-1^) and the C18:1-to-C18:0 ratio. **(M)** Elongation index [(C18:0+C18:1)/C16:0] of fatty acyl chains in liver PLs. n = 6-7 per group. Data are mean ± SEM. *p<0.05, **p<0.01, ***p<0.001, and ****p<0.0001 by two-way ANOVA followed by Tukey’s post hoc tests.

In addition to total concentration, we determined the composition of fatty acyl chains in glycerolipids. The composition of all DAG fatty acyl chains was comparable between untrained WT and LAKO mice (Fig. 4E-G). Training decreased C16:1, C18:1, and C18:2 in liver DAGs of WT mice (Fig. 4E and F). Thus, training led to a more saturated fatty acyl composition as exemplified by the lower desaturation indexes (C16:1-to-C16:0 and C18:1-to-18:0 ratios) of liver DAGs in trained WT mice (Fig. 4E). In contrast, training did not change C16:1, and the C16:1-to-C16:0 ratio in LAKO mice (Fig. 4E). Training did decrease C18:1, C18:2, and the C18:1-to-C18:0 in liver DAGs of LAKO mice (Fig. 4F), however, the training response was less pronounced when compared to WT mice (Fig. 4F). The elongation index [(C18:0+C18:1)/C16:0] was reduced in trained WT mice, but not trained LAKO mice (Fig. 4G). It should be noted that livers of WT-trained mice also showed lower unsaturated and elongated fatty acids comprising DAGs compared to LAKO-trained mice following an acute exercise bout (Supplemental Fig. S1) supporting that these changes in DAGs were the result of training adaptations and not an acute response to the last exercise bout. Together, these results indicate that training lowers the concentration of unsaturated and elongated of fatty acids comprising liver DAGs in an AMPK-dependent manner.

Similar to DAGs, the composition of liver TAG fatty acyl chains was comparable between untrained WT and LAKO mice (Fig. 4H-J). In contrast to DAGs, training did not lower unsaturated fatty acids comprising liver TAGs in WT mice (Fig. 4H-J). Training lowered C18:0 and thereby increased the desaturation index (C18:1-to-C18:0 ratio) of fatty acids in liver TAGs of WT mice (Fig. 4I). Interestingly, training increased the monounsaturated fatty acids of liver TAGs in LAKO mice as indicated by higher C16:1 and C18:1 (Fig. 4H and I). Training also increased the elongation index of fatty acids comprising liver TAGs in LAKO mice (Fig. 4J). These results indicate that a significant training-induced decline in unsaturated and elongated fatty acids was not observed in liver TAGs as it was in DAGs. However, liver TAG accretion as well as the desaturation and elongation of fatty acyl chains comprising TAGs was increased in exercise-trained LAKO mice.

Changes in phospholipid fatty acyl chain composition were modest compared to those seen in either DAGs or TAGs. Training decreased the concentration of C16:0 and C18:2 fatty acyl chains in liver phospholipids of WT mice (Fig. 4K-M). In LAKO mice, training increased C16:1 in liver phospholipids (Fig. 4K). Together, our results suggest that AMPK regulates both the concentration and composition of liver lipids in response to training.

### Hepatic AMPK is necessary for glycogen deposition in response to exercise training

The role of AMPK on glucose metabolism was tested in response to training. Liver glycogen was comparable between genotypes in untrained mice (Fig. 5A). Training increased liver glycogen in WT mice by ∼3-fold (Fig. 5A). This training-induced expansion of the glycogen pool did not occur in LAKO mice (Fig. 5A). Molecular regulators of glycogen metabolism were also assessed. Liver glycogen phosphorylase (PYGL) (Fig. 5B and F) and the phospho-glycogen synthase (pGS)-to-GS ratio (Fig. 5C and F) did not differ between genotypes or in response to training. Regulators of glucose-6-phosphate, a glycogen precursor/product, were determined. A genotype effect showed liver glucose-6-phosphatase (G6PC) protein was higher in the absence of AMPK (Fig. 5D and F), while phosphoenolpyruvate carboxykinase 1 (PCK1) protein was unaffected (Fig. 5E and F).

**FIGURE 5.**
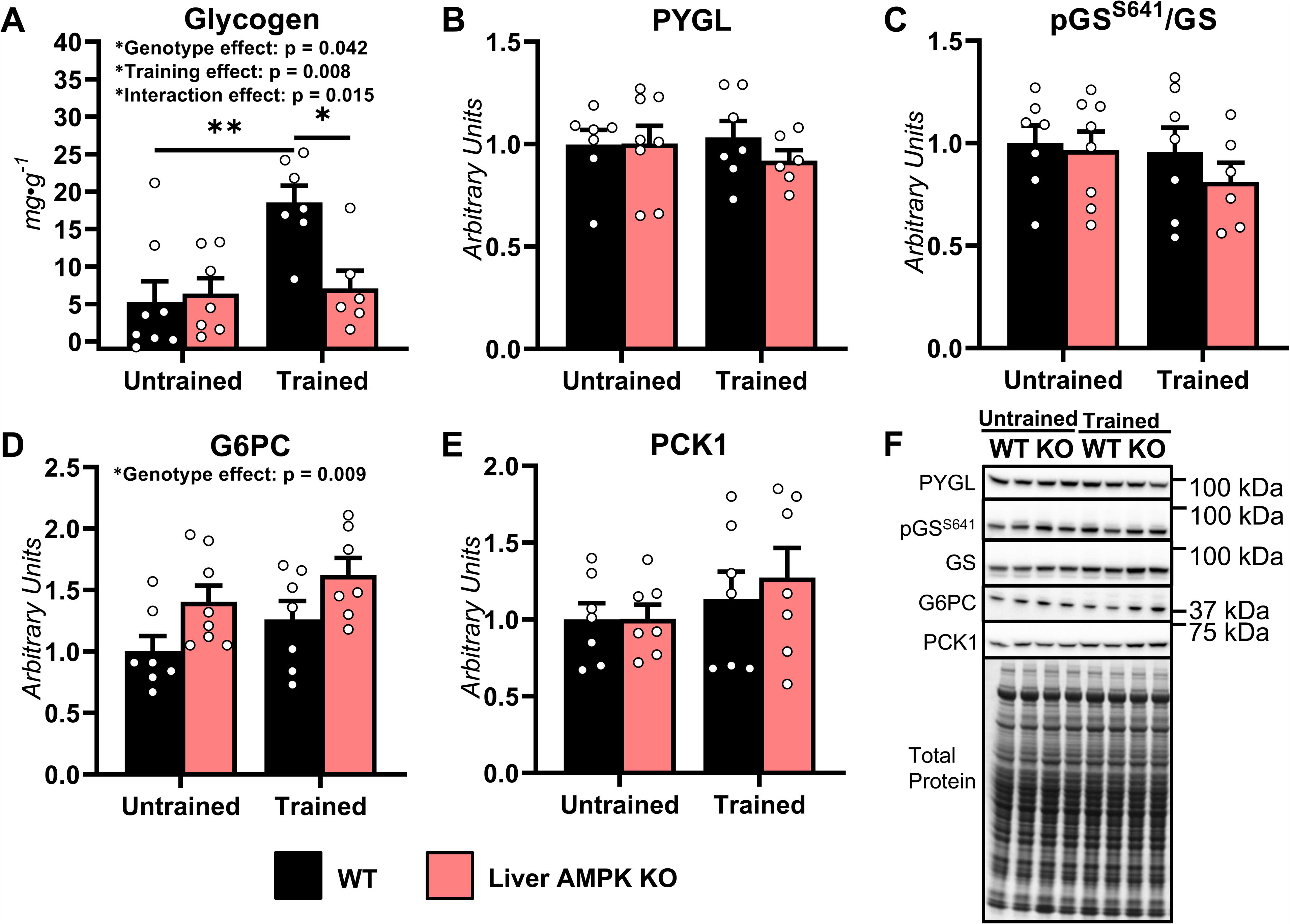
Glycogen and molecular regulators of glycogen homeostasis in untrained and exercise-trained mice lacking hepatic AMPK. Livers from untrained and trained mice with liver-specific deletions of AMPK α1 and α2 subunits (Liver AMPK KO) and wild type (WT) littermates were obtained from non-catheterized mice on a stationary treadmill at the 0-minute time point (8.5-hour fast). **(A)** Liver glycogen content (mg·g^-1^). Liver **(B)** glycogen phosphorylase (PYGL), **(C)** phosphorylated glycogen synthase-to-total glycogen synthase ratio (pGS^S641^/GS), **(D)** glucose-6-phosphatase, catalytic subunit (G6PC), and **(E)** cytosolic phosphoenolpyruvate carboxykinase (PCK1) as determined by immunoblotting. **(F)** Representative immunoblots for liver PYGL, pGS^S641^, total GS, G6PC, and PCK1. n = 6-8 per group. Data are mean ± SEM. *p<0.05 and **p<0.01 by two-way ANOVA followed by Tukey’s post hoc tests.

### Glycogenolysis, but not gluconeogenesis and mitochondrial oxidative fluxes, is blunted during acute exercise in trained mice lacking hepatic AMPK

Glycogen is a key source of liver glucose production during exercise. As such, glucose production and associated nutrient fluxes (Fig. 6A-N) were quantified during acute exercise in trained mice to test the impact of lower glycogen levels in LAKO mice on glucose homeostasis during an exercise bout. Arterial glucose was comparable between trained WT and LAKO mice at rest (Fig. 6B). In contrast, circulating glucose was lower in LAKO mice compared to WT mice at 10 minutes of exercise and trended lower throughout the remainder of the 30-minute exercise bout (Fig 6B). Similarly, endogenous glucose production (V_EGP_) was comparable between genotypes at rest and lower in LAKO mice compared to WT mice during acute exercise (Fig. 6C). The blunted rise in endogenous glucose production during exercise in the LAKO mice was accompanied by diminished glycogenolysis (V_PYGL_) (Fig. 6D). Total gluconeogenesis (V_Aldo_) increased during muscular work in both WT and LAKO mice (Fig. 6E) and this was paralleled by elevated rates of gluconeogenesis from phosphoenolpyruvate (V_Enol_; Fig. 6G) and tricarboxylic acid (TCA) cycle cataplerosis (V_PCK_; Fig. 6H). Anaplerosis and related fluxes were higher in response to exercise in both WT and LAKO mice (Fig. 6I-K). This included the flux of non-phosphoenolpyruvate anaplerotic sources to pyruvate (V_LDH_; Fig. 6I), flux of pyruvate to oxaloacetate (V_PC_; Fig. 6J), and flux from propionyl-CoA to succinyl-CoA (V_PCC_; Fig. 6K). TCA cycle fluxes (V_CS_ and V_SDH_) and pyruvate cycling (V_PK+ME_) increased similarly in both genotypes during muscular work (Fig. 6L-M). These results indicate that the inability of training to increase liver glycogen stores in LAKO mice compromises the capacity for glycogenolysis and glucose production during subsequent acute exercise bouts.

**FIGURE 6.**
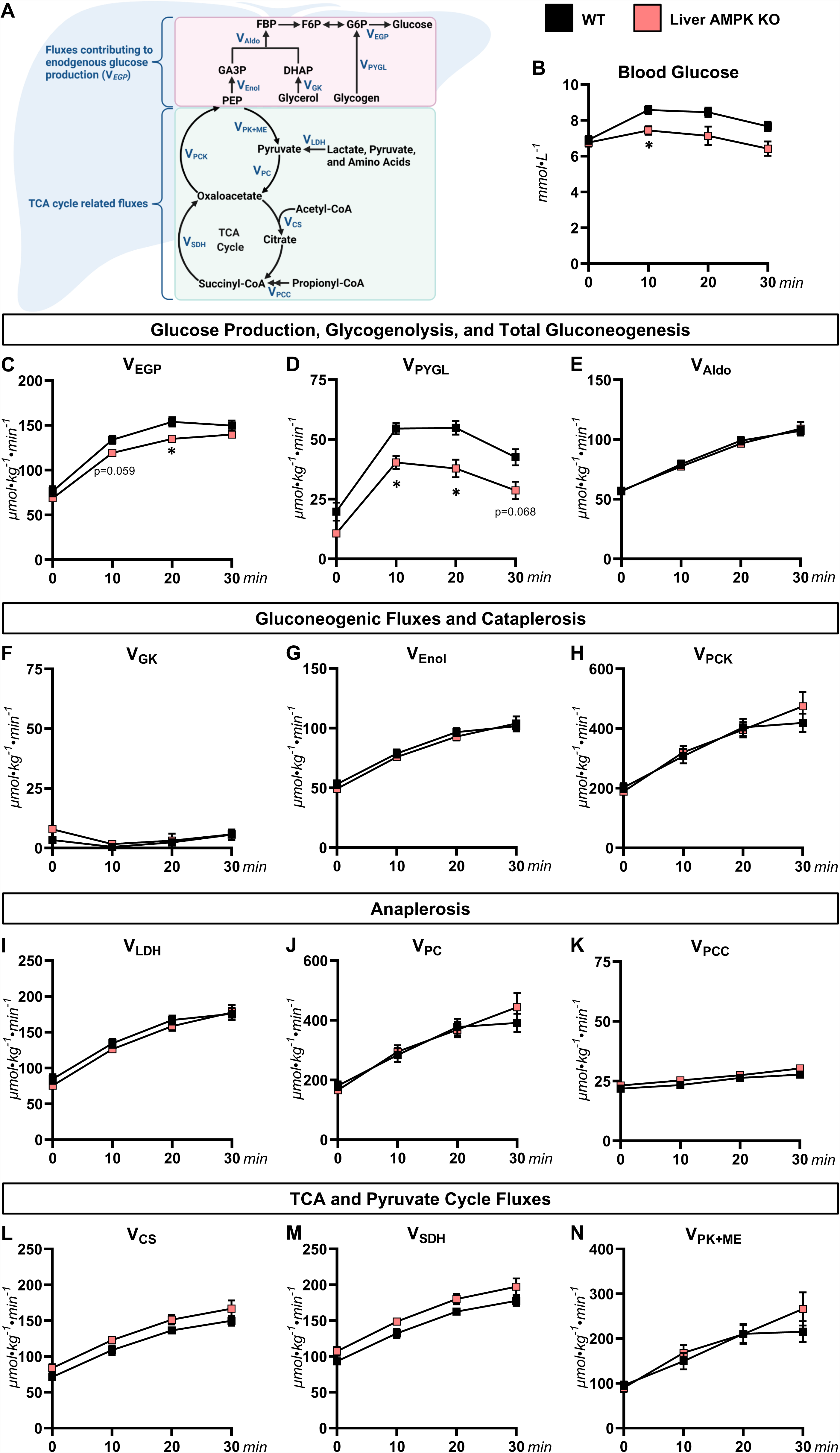
Glucose producing and oxidative metabolism fluxes prior to and during treadmill running in exercise-trained mice lacking liver AMPK. **(A)** A schematic of model-estimated glucose producing and oxidative fluxes. **(B)** A time course of arterial blood glucose concentration in mice with liver-specific deletions of AMPK α1 and α2 subunits (Liver AMPK KO) and wild type (WT) littermates prior to and during a 30-minute treadmill run. Model-estimated, absolute nutrient fluxes (μmol·kg^-1^·min^-1^) in WT and KO mice prior to and during a 30-minute of treadmill run for **(C)** endogenous glucose production (V_EGP_), **(D)** glycogenolysis (V_PYGL_), **(E)** total gluconeogenesis (V_Aldo_), **(F)** gluconeogenesis from glycerol (V_GK_), **(G)** gluconeogenesis from phosphoenolpyruvate (V_Enol_), **(H)** tricarboxylic acid cycle cataplerosis (V_PCK_), **(I)** flux from unlabeled, non-phosphoenolpyruvate, anaplerotic sources to pyruvate (V_LDH_), **(J)** anaplerosis from pyruvate (V_PC_), **(K)** anaplerosis from propionyl-CoA (V_PCC_), **(L)** flux from oxaloacetate and acetyl-CoA to citrate (V_CS_), **(M)** flux from succinyl-CoA to oxaloacetate (V_SDH_), and **(N)** pyruvate cycling (V*_PK+ME_*). n = 5-8 per genotype. Data are mean ± SEM. *p<0.05 vs. WT at specified time point by two-way repeated-measures ANOVA followed by Šidák’s post hoc tests.

### Differential impact of liver AMPK during acute exercise on glycogen and glycerolipids

Liver metabolites were measured in trained mice following acute exercise and subtracted from resting controls to assess net changes in response to a bout of exercise (Fig. 7A-H). Net changes in and absolute concentrations of liver adenine nucleotides and energy charge following acute exercise were similar between WT and LAKO mice (Fig. 7A-D). The decline in liver glycogen was attenuated in LAKO mice compared to WT mice (Fig. 7E). Glycogen levels at the end of the acute exercise bout were lower in LAKO mice (Fig. 7E). This is likely due to less liver glycogen in LAKO mice prior to initiating the exercise bout (Fig. 5A). Interestingly, LAKO mice showed a more pronounced decrease in liver DAGs during exercise. However, liver DAG levels remained higher in LAKO mice at the end of the exercise bout (Fig. 7F). Exercise reduced liver TAG in LAKO mice, but not WT mice (Fig. 7G). This resulted in the normalization of liver TAGs between genotypes after acute exercise (Fig. 7G).

**FIGURE 7.**
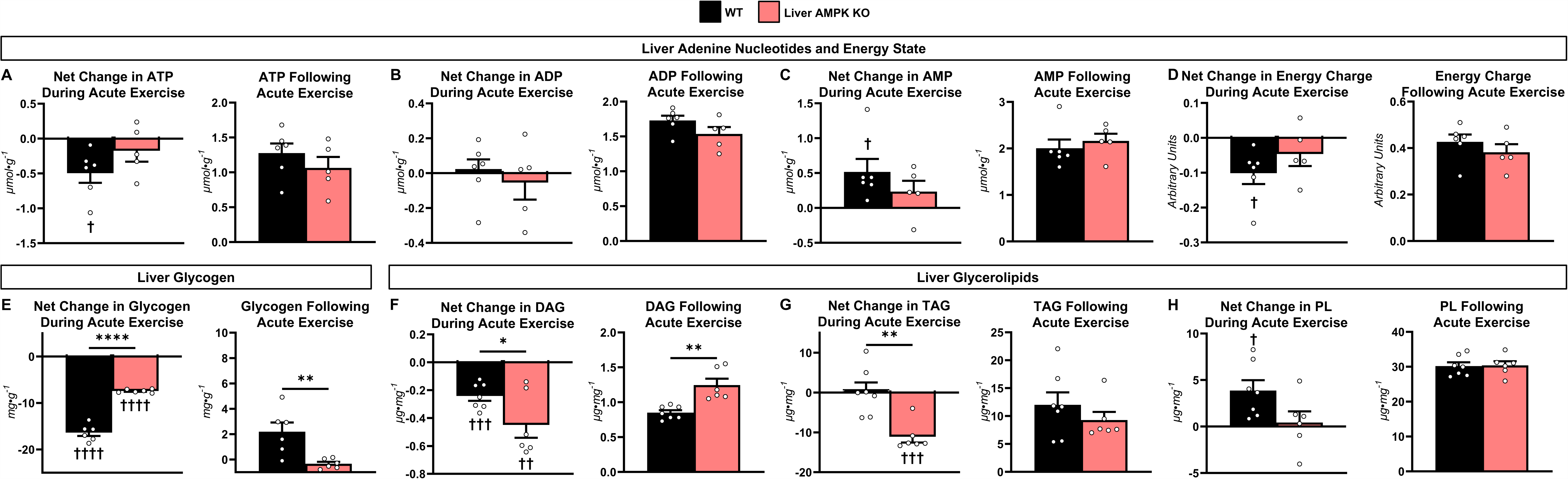
Impact of an acute exercise bout on liver metabolites in exercise-trained mice lacking liver AMPK. Livers metabolites from mice exhibiting liver-specific deletions of AMPK α1 and α2 subunits (Liver AMPK KO) and wild type mice (WT) littermates. **(A)** Net Change in liver ATP during acute exercise and total liver ATP following acute exercise (µmol·g^-1^). **(B)** Net change in liver ADP during acute exercise and total liver ADP following acute exercise (µmol·g^-1^). **(C)** Net change in liver AMP during acute exercise and total liver AMP following acute exercise (µmol·g^-1^). **(D)** Net change in liver energy charge during acute exercise (arbitrary units; [ATP+0.5ADP]/[TAN]) and liver energy charge following acute exercise. **(E)** Net change in liver glycogen during acute exercise and total liver glycogen following acute exercise (mg·g^-1^). **(F)** Net change in liver diacylglycerides (DAGs) during acute exercise and total liver DAGs following acute exercise (µg·mg^-1^). **(G)** Net change in liver triacylglycerides (TAGs) during acute exercise and total liver TAGs following acute exercise (µg·mg^-1^). **(H)** Net change in liver phospholipids (PLs) during acute exercise and total liver PLs following acute exercise (µg·mg^-1^). n = 5-7 per genotype. Data are mean ± SEM. *p<0.05, **p<0.01, and ****p<0.0001 between genotypes determined by Students *t* test. Change in metabolites were calculated as the difference between rest (mean concentration) and post-exercise concentrations. Note: Liver metabolites measured at rest and post-exercise were from non-catheterized and catheterized mice, respectively. †p<0.05, ††p<0.01, †††p<0.001, and ††††p<0.0001 by one sample t tests using theoretical mean of zero to determine impact of exercise within genotype.

## DISCUSSION

Exercise training elicits adaptations in liver metabolism that facilitate energy provision in subsequent exercise bouts and mitigate the metabolic dysregulation characterizing chronic diseases such as non-alcoholic fatty liver disease and insulin resistance (8, 9). A decline in liver energy state is a fundamental response to acute exercise (Fig. 7A-C) (5, 23). However, whether repeated hepatic energetic discharge elicited during training is responsible for the metabolic adaptations to training in the liver and the underlying molecular mechanisms of action remain to be clearly defined. In this study we tested the hypothesis that the energy sensor and effector, AMPK, is necessary for liver glucose and lipid adaptations to exercise training. Novel findings of this study were that training is dependent on hepatic AMPK to promote i) higher liver glycogen that allows for accelerated glycogenolysis and glucose production during acute exercise, ii) a decrease in liver glycerolipids independent of TCA cycle flux, and iii) lower desaturation and elongation of fatty acids in liver DAGs.

### Role of AMPK in glucose adaptations to exercise training

Our prior work showed that deletion of hepatic AMPKα1α2 does not impact blood glucose concentration, endogenous glucose production, and gluconeogenesis in fasted, sedentary mice (5, 26, 34, 38, 39). However, it should be noted that independent groups have hypothesized that hepatic AMPK activation impedes the release of glucose from the liver (40). This is based on studies where liver PCK1 and/or G6Pase mRNA as well as blood glucose are reduced in mice with constitutively active AMPKα1 or α2 subunits (41, 42). Further, it was shown that mutating AMPKγ1 to constitutively activate AMPK suppressed glucose output in hepatocytes (43) and deletion of hepatic AMPKα2 elevated glucose production in fasted mice (44). The hypothesis that AMPK is an inhibitor of liver glucose production is difficult to reconcile with physiology as AMPK activity and endogenous glucose production both increase during a condition such as acute exercise (5, 23). The discrepancy may be due to the approach of the present study and our prior work where both α subunits of AMPK are deleted. Research that suggested AMPK inhibits endogenous glucose production was done using either the deletion of a single α subunit (44) or approaches that used constitutively activate AMPK (41–43). The experiments presented here tested the impact of repeated activation of AMPK as a physiological response to single exercise bouts (23). Liver G6Pase protein was slightly increased in LAKO mice. However, circulating glucose and glucose fluxes (endogenous glucose production, glycogenolysis, and gluconeogenesis) were similar between trained WT and LAKO mice under fasted, resting conditions. Together with prior work (5, 26, 34, 38, 39), these results support that AMPK does not inhibit liver glucose production under resting conditions in the absence of an acute provocative stimulus.

While resting glucose fluxes are not impacted by loss of AMPK, mice with whole-body and liver-specific disruptions in AMPK have lower liver glycogen concentrations under resting conditions (5, 45, 46). Our previous work also determined that refeeding following a fast does not replenish liver glycogen to as high a concentration in mice lacking hepatic AMPKα1α2 as that observed in WT mice (5). This suggests AMPK is necessary for a robust deposition of liver glycogen. Exercise training in rodent models usually (10–16), but not always (47–49), causes increased resting liver glycogen. Here we show six weeks of training elevates liver glycogen by ∼3-fold in WT mice in the postabsorptive, sedentary state. Remarkably, this increase in liver glycogen was completely attenuated in LAKO mice. Our data in the liver is consistent with and extends previous experiments in mice with a skeletal muscle-specific deletion of AMPKα1α2 showing that AMPK is necessary for increased muscle glycogen synthesis in the recovery period following acute exercise (50). Since glycogen is an important source of liver glucose production during acute exercise, we quantified glucose fluxes in mice during a 30-minute treadmill run.

Trained WT mice had higher arterial blood glucose and endogenous glucose production compared to trained LAKO mice during an acute exercise challenge. The heightened rate of glucose production in trained WT mice was linked to higher glycogenolysis compared to trained LAKO mice. Liver glycogen concentration has a positive relationship with time to exhaustion during acute exercise in both mice and humans (51, 52). Exercise capacity as determined by maximal running speed during an exercise stress test increased in both WT and LAKO mice following exercise training. An exercise stress test was performed in this study, rather than exercise endurance protocol, because it is generally assumed to be cardiopulmonary-limited given that it positively correlates with maximum whole body oxygen uptake (24). The design of the exercise stress test is a short challenge and is not likely limited by the difference in nutrient availability which may occur in LAKO mice. This allowed the documentation of a training effect without potential limitations due to lower liver glycogen in LAKO mice. Nevertheless, by promoting deposition of glycogen in response to exercise training, AMPK allows for accelerated glycogenolysis and glucose production during subsequent acute exercise and facilitates glucose supply to working muscle.

### Role of AMPK in adaptations of hepatic lipids to exercise training

In addition to control of glucose production, we assessed liver lipid concentration and composition in untrained and trained WT and LAKO mice. Regular exercise is often considered to be a potential means of improving the lipid liver profile. The key mechanisms for any improvement remain to be fully defined. Liver DAGs and TAGs in untrained WT mice were comparable to untrained LAKO mice under fasted, resting conditions. This is consistent with prior studies showing unchanged (5, 26, 34, 38, 53, 54) more often than increased (5, 53, 55) liver DAGs and TAGs in sedentary mice with disruptions in AMPK. We previously reported that liver TAGs were similar in liver-specific AMPKα1α2 KO mice compared to WT littermates under sedentary conditions (34). In those same studies we showed that AMPKα1α2 is necessary for the decline in liver TAGs following acute administration of AMP mimetic, 5-aminoimidazole-4-carboxamide-1-β-D-ribofuranoside (34). This suggests that conditions provoking energetic discharge unmask otherwise silent lipid phenotypes mediated by AMPK. In agreement with this notion, the use of pharmacological AMPK activators in AMPK knockout models and constitutively active AMPK models consistently show that AMPK lowers liver DAGs and TAGs (38, 42, 43, 56, 57). In this study we tested the hypothesis that hepatic AMPK is necessary for the repeated decrease in liver energy state that occurs with training to lower liver lipids. Training decreased liver DAGs in both genotypes. However, the decline in DAGs was attenuated in LAKO mice. Moreover, training led to higher liver TAGs in LAKO mice. Thus, our results indicate that AMPK is necessary for the full liver lipid lowering effects of training.

AMPK promotes mitochondrial biogenesis and positively regulates mitochondrial oxidative metabolism (i.e., β-oxidation, TCA cycle flux, oxidative phosphorylation, and ketogenesis) (58). Given this, we assessed mitochondrial pathways involved in lipid disposal in WT and LAKO mice. In contrast to resting conditions, acute exercise decreased liver TAGs in trained LAKO mice to levels observed in trained WT mice. Acute exercise also diminished liver DAGs in trained LAKO mice, but they remained higher than DAGs in livers of trained WT mice. The persistent elevation in liver DAGs following both acute exercise and exercise training could be due to impaired lipid disposal in LAKO mice. We previously determined that mitochondrial respiratory function supported by NADH-linked substrates was impaired by loss of hepatic AMPK (34). Here we observed lower mitochondrial respiratory chain proteins (complexes I, III, and IV) in LAKO mice compared WT mice. However, TCA cycle flux was comparable between genotypes at rest and during acute exercise suggesting lipid oxidation to CO_2_ in the liver is not diminished by deletion of AMPK *in vivo*. Prior work in which hepatocytes or liver homogenate was incubated with ^14^C-labeled palmitate resulted in an AMPK-dependent increase in ^14^C-labeled acid soluble products, but not ^14^CO_2_ combined with ^14^C-labeled acid soluble products (38, 43, 56). In addition, AMPK activation increases ketone bodies (41, 56). Thus, the reduction of liver lipids following training may be mediated through an AMPK-dependent increment in β-oxidation and/or ketogenesis and not by accelerated TCA cycle flux. Notably, we observed that the net decrease in liver DAGs and TAGs following acute exercise were more pronounced in LAKO mice. Given this, further studies are necessary to define the significance of lipid disposal pathways in mediating the elevated liver lipids in LAKO mice following training.

The inhibition of lipid synthesis and storage are additional functions attributed to AMPK (58). A mechanism through which AMPK suppresses fatty acid synthesis is the reversible phosphorylation of ACC (58). We observed that ACC phosphorylation was abolished in both untrained and trained LAKO mice. Importantly, Marcinko et al. determined that mice with whole body knock-in mutations of AMPK phosphorylation sites on ACC, which results in constitutively active ACC, do not exhibit dysregulated liver lipid accretion following exercise training (59). This suggests that other mechanisms are responsible for elevated glycerolipids in trained LAKO mice. An often reported finding in rodent studies is the reduction of liver MUFAs in response to exercise training (60). This exercise-induced lowering of MUFAs is linked to decreased activity of SCD1, which catalyzes the synthesis of MUFAs that are preferentially incorporated into glycerolipids (60, 61). In agreement, we observed an overt decline in the concentration of MUFAs comprising liver DAGs in WT mice following training that was not apparent in LAKO mice. Moreover, liver SCD1 protein expression was higher in LAKO mice. Hepatic AMPK phosphorylates and inhibits sterol regulatory element binding protein-1c (SREBP-1c) (62, 63). Thus, exercise training may limit liver lipid accretion by promoting AMPK translocation to the endoplasmic reticulum where it inhibits SREBP-1c and, subsequently, SCD-1 expression. This would limit fatty acid desaturation and their preferential incorporation into glycerolipids. This may also be a significant mechanism through which exercise prevents NAFLD, which is commonly characterized by an increase in hepatic MUFAs (64).

It should also be noted that training decreased the percentage of MUFAs comprising liver DAGs but not TAGs in WT mice. SCD1 co-localizes with diacylglycerol-O-acyltransferase (DGAT) 2, which catalyzes the final step of TAG synthesis, at the endoplasmic reticulum (65). Prior work showed this close spatial proximity between SCD1 and DGAT2 promotes the use of MUFAs in triglyceride synthesis (65). In addition, DGAT1 has been speculated to prefer MUFAs as a substrate over saturated fatty acids (66). Given that the liver DAG pool size is considerably smaller than that of TAGs, preferential channeling of MUFAs into TAGs (via DGAT localization and/or activity) could mask the training-induced lowering of unsaturated fatty acids comprising DAGs.

In conclusion, AMPK is necessary for adaptations to training in liver glycogen dynamics that allow the liver to effectively meet glucose demands of working muscle by increasing the capacity for glycogenolysis. AMPK also mediates the inhibition of lipid accretion by training in a manner linked to lowering the desaturation and elongation of fatty acids comprising DAGs. Thus, AMPK is required for liver training adaptations that are critical to glucose and lipid metabolism at rest and when confronted with an exercise challenge. These studies provide a mechanistic foundation for regular exercise as a means of preventing and possibly reversing fatty liver.

## ACKNOWLEDGEMENTS

The authors acknowledge the technical assistance provided by the Vanderbilt University Mouse Metabolic Phenotyping Center Analytical Core Services. Figure schematics were created using BioRender.com.

## AUTHOR CONTRIBUTIONS

CCH, MF, BV, and DHW conceived and designed the experiments. CCH, DPB, FIR, MG, and EPD performed experiments. CCH analyzed and interpreted data. CCH drafted the manuscript. CCH, DHW, MF, and BV edited and revised the manuscript. All authors approved the final version of the manuscript for publication.

---

**SUPPLEMENTAL FIGURE S1.**
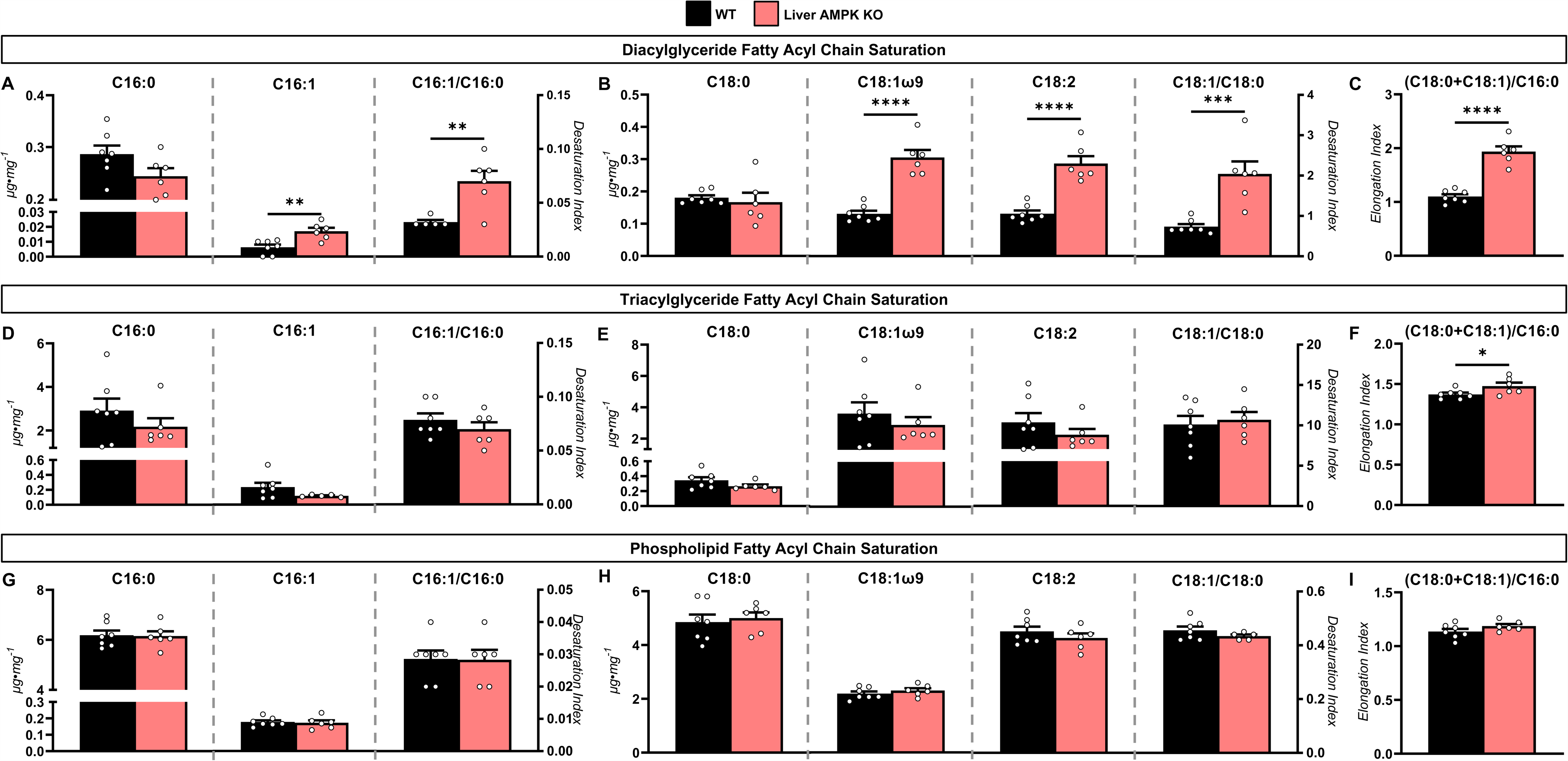
Fatty acyl chain composition comprising glycerolipids of exercise-trained mice following an acute exercise bout. Livers from trained mice exhibiting liver-specific deletions of AMPK α1 and α2 subunits (Liver AMPK KO) and wild type (WT) littermates were obtained from catheterized mice following a 30-minute treadmill running bout. **(A)** C16:0 and C16:1 fatty acids in liver diacylglycerides (µg·mg^-1^; DAGs) and the C16:1-to-C16:0 ratio. **(B)** C18:0, C18:1, and C18:2 fatty acids in liver DAGs (µg·mg^-1^) and the C18:1-to-C18:0 ratio. **(C)** Elongation index [(C18:0+C18:1)/C16:0] of fatty acyl chains in liver DAGs. **(D)** C16:0 and C16:1 fatty acids in liver triacylglycerides (µg·mg^-1^; TAGs) and the C16:1-to-C16:0 ratio. **(E)** C18:0, C18:1, and 18:2 fatty acids in liver TAGs and the C18:1-to-C18:0 ratio. **(F)** Elongation index [(C18:0+C18:1)/C16:0] of fatty acyl chains in liver TAGs. **(G)** C16:0 and C16:1 fatty acids in liver phospholipids (µg·mg^-1^; PLs) and the C16:1-to-C16:0 ratio. **(H)** C18:0, C18:1, and C18:2 fatty acids in liver PLs (µg·mg^-1^) and the C18:1-to-C18:0 ratio. **(I)** Elongation index [(C18:0+C18:1)/C16:0] of fatty acyl chains in liver PLs. n = 5-7 per group. Data are mean ± SEM. *p<0.05, **p<0.01, ***p<0.001, and ****p<0.0001 by Students *t* tests.

**Supplemental Table S1.**
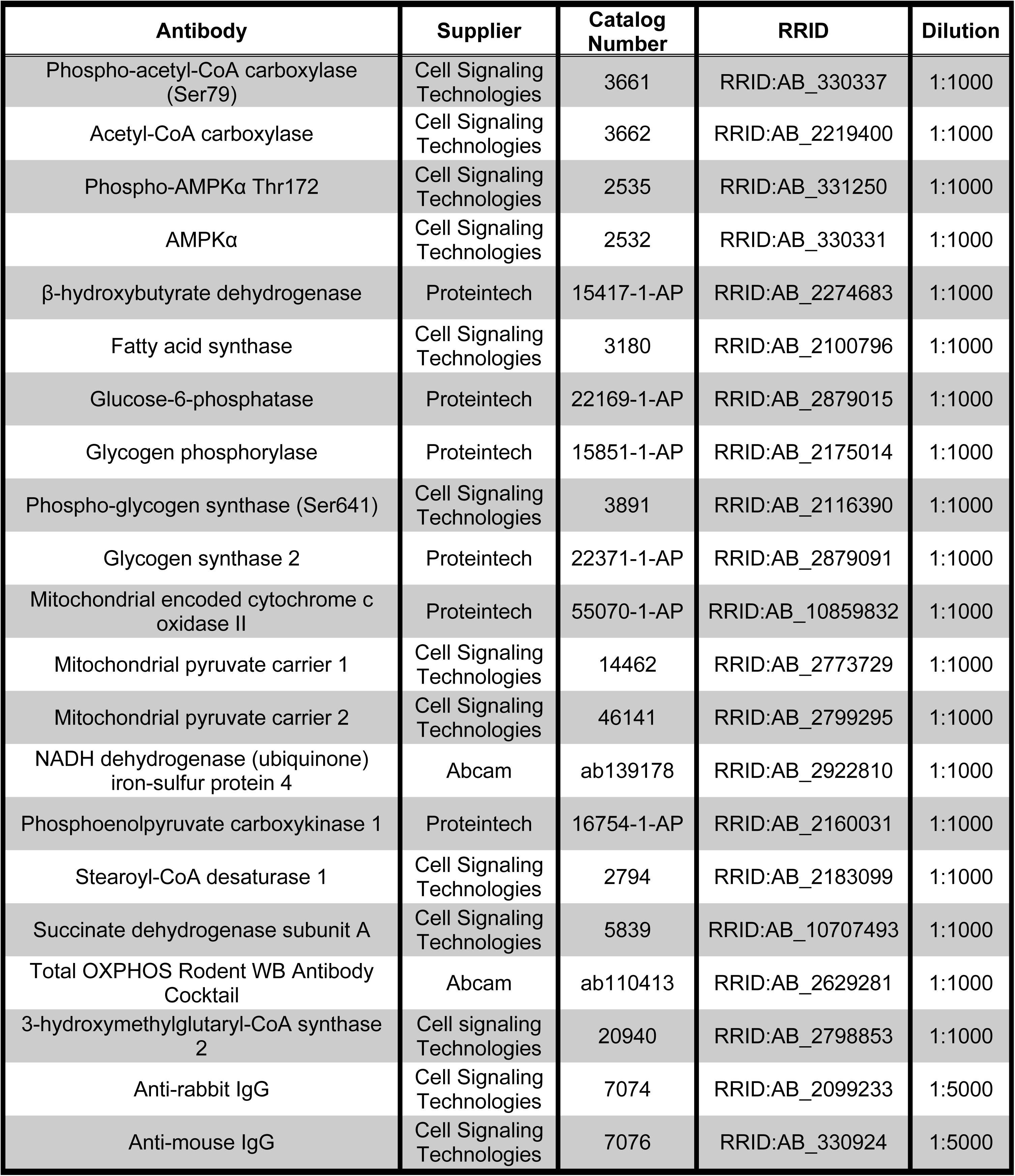
List of antibodies.

| Antibody | Supplier | Catalog Number | RRID | Dilution |
| --- | --- | --- | --- | --- |
| Phospho-acetyl-CoA carboxylase (Ser79) | Cell Signaling Technologies | 3661 | RRID:AB_330337 | 1:1000 |
| Acetyl-CoA carboxylase | Cell Signaling Technologies | 3662 | RRID:AB_2219400 | 1:1000 |
| Phospho-AMPKα Thr172 | Cell Signaling Technologies | 2535 | RRID:AB_331250 | 1:1000 |
| AMPKα | Cell Signaling Technologies | 2532 | RRID:AB_330331 | 1:1000 |
| β-hydroxybutyrate dehydrogenase | Proteintech | 15417-1-AP | RRID:AB_2274683 | 1:1000 |
| Fatty acid synthase | Cell Signaling Technologies | 3180 | RRID:AB_2100796 | 1:1000 |
| Glucose-6-phosphatase | Proteintech | 22169-1-AP | RRID:AB_2879015 | 1:1000 |
| Glycogen phosphorylase | Proteintech | 15851-1-AP | RRID:AB_2175014 | 1:1000 |
| Phospho-glycogen synthase (Ser641) | Cell Signaling Technologies | 3891 | RRID:AB_2116390 | 1:1000 |
| Glycogen synthase 2 | Proteintech | 22371-1-AP | RRID:AB_2879091 | 1:1000 |
| Mitochondrial encoded cytochrome c oxidase II | Proteintech | 55070-1-AP | RRID:AB_10859832 | 1:1000 |
| Mitochondrial pyruvate carrier 1 | Cell Signaling Technologies | 14462 | RRID:AB_2773729 | 1:1000 |
| Mitochondrial pyruvate carrier 2 | Cell Signaling Technologies | 46141 | RRID:AB_2799295 | 1:1000 |
| NADH dehydrogenase (ubiquinone) iron-sulfur protein 4 | Abcam | ab139178 | RRID:AB_2922810 | 1:1000 |
| Phosphoenolpyruvate carboxykinase 1 | Proteintech | 16754-1-AP | RRID:AB_2160031 | 1:1000 |
| Stearoyl-CoA desaturase 1 | Cell Signaling Technologies | 2794 | RRID:AB_2183099 | 1:1000 |
| Succinate dehydrogenase subunit A | Cell Signaling Technologies | 5839 | RRID:AB_10707493 | 1:1000 |
| Total OXPHOS Rodent WB Antibody Cocktail | Abcam | ab110413 | RRID:AB_2629281 | 1:1000 |
| 3-hydroxymethylglutaryl-CoA synthase 2 | Cell signaling Technologies | 20940 | RRID:AB_2798853 | 1:1000 |
| Anti-rabbit IgG | Cell Signaling Technologies | 7074 | RRID:AB_2099233 | 1:5000 |
| Anti-mouse IgG | Cell Signaling Technologies | 7076 | RRID:AB_330924 | 1:5000 |

